# Reliability of Decision-Making and Reinforcement Learning Computational Parameters

**DOI:** 10.1101/2021.06.30.450026

**Authors:** Anahit Mkrtchian, Vincent Valton, Jonathan P. Roiser

## Abstract

Computational models can offer mechanistic insight into cognition and therefore have the potential to transform our understanding of psychiatric disorders and their treatment. For translational efforts to be successful, it is imperative that computational measures capture individual characteristics reliably. Here we examine the reliability of reinforcement learning and economic models derived from two commonly used tasks. Healthy individuals (N=50) completed a restless four-armed bandit and a calibrated gambling task twice, two weeks apart. Reward and punishment learning rates from the reinforcement learning model showed good reliability and reward and punishment sensitivity from the same model had fair reliability; while risk aversion and loss aversion parameters from a prospect theory model exhibited good and excellent reliability, respectively. Both models were further able to predict future behaviour above chance within individuals. This prediction was better when based on participants’ own model parameters than other participants’ parameter estimates. These results suggest that reinforcement learning, and particularly prospect theory parameters, as derived from a restless four-armed bandit and a calibrated gambling task, can be measured reliably to assess learning and decision-making mechanisms. Overall, these findings indicate the translational potential of clinically-relevant computational parameters for precision psychiatry.

## Introduction

Cognitive and neural processes are increasingly conceptualized in computational terms (Palminteri et al., 2017). Generative computational models offer the advantage of examining behaviourally unobservable, but important, latent processes that drive behaviour and can be closely linked to neurobiology (Huys et al., 2016; Montague et al., 2004). As such, they provide a mathematically precise framework for specifying hypotheses about the cognitive processes that generate behaviour. These features make modelling a powerful tool to provide mechanistic accounts into (a)typical behaviours, including those associated with psychiatric symptoms (Browning et al., 2020; Montague et al., 2012).

One area of computational modelling that has received particular attention is the reward and punishment processes underlying decision-making (Maia, 2009). In particular, two classes of computational models, respectively originating from computer science and behavioural economics, have been influential in characterizing the cognitive mechanisms underlying decision-making: reinforcement learning (RL) and prospect theory (PT) (Dayan & Niv, 2008; Kahneman & Tversky, 1979; Maia, 2009; Niv, 2009; Schonberg et al., 2011; Sutton & Barto, 2018; Tversky & Kahneman, 1992). RL models describe how agents learn from rewards and punishments through trial-and-error (Sutton & Barto, 2018). Within the field of computational psychiatry – which aims to better understand psychiatric symptoms through computational methods – this set of models has perhaps been the most influential. For example, reward and punishment sensitivity (reflecting subjective valuation of the outcomes) and learning rate (reflecting how quickly individuals learn from better- or worse-than-expected outcomes) have been associated with distinct symptomatology and neural signals (Daw & Doya, 2006; Huys et al., 2021; Maia & Frank, 2011; Niv, 2009). A commonly-used RL task is the multi-armed “bandit”. On this task individuals choose between multiple slot machines with fluctuating, unknown probabilities of reward and punishment, with the goal of maximizing earnings (Daw et al., 2006; Seymour et al., 2012; Speekenbrink & Konstantinidis, 2015; Yi et al., 2009). Participants must decide on each trial whether to persist with the previously sampled slot machine or explore others which may yield better outcomes. The mechanisms thought to underlie these decisions can be captured effectively by RL models (Aylward et al., 2019; Daw et al., 2006).

PT models, on the other hand, describe the cognitive processes driving decision-making biases under known risks, and have been extremely influential in understanding economic decision-making (Kahneman & Tversky, 1979; Ruggeri et al., 2020; Schonberg et al., 2011; Sokol-Hessner & Rutledge, 2019; Tversky & Kahneman, 1992). These processes are often examined by asking participants to choose between a guaranteed outcome (e.g., £0 gain) and a 50% gamble with two possible outcomes (e.g., £30 gain or £10 loss). It is commonly observed that humans tend to prefer a sure payment over a risky payment with equivalent or higher expected value. For example, you may prefer an investment with a fixed return over one with a potentially higher but uncertain return. PT proposes that these observations can be accounted for by two different processes: 1) risk aversion – the preference for certain over uncertain gains, and 2) loss aversion – weighting losses more heavily than gains. Risk and loss aversion vary across individuals, and these differences have been associated with various psychiatric states. For example, greater risk aversion has been observed in anxiety, while loss aversion has been associated with obsessive-compulsive disorder as well as suicidality (Baek et al., 2017; Brown et al., 2013; Charpentier et al., 2017; Charpentier et al., 2016; Hadlaczky et al., 2018; Hartley & Phelps, 2012; Klaus et al., 2020; Sip et al., 2017; Stauffer et al., 2014; Tobler et al., 2009; Tremeau et al., 2008). Importantly, computational modelling has allowed researchers to dissociate risk and loss aversion and their contribution to symptoms (Charpentier et al., 2017).

Parameters from RL and PT models thus show promise in generating insights into the mechanisms underlying psychiatric symptoms. For such translational endeavours to be successful, however, it is vital that computational measures capture individual characteristics reliably (Browning et al., 2020; Paulus et al., 2016). Specifically, the reliability of measures set an upper limit for detecting both relationships with other measures, e.g., symptoms, and the effect of treatment interventions in e.g., randomized controlled trials, which may otherwise be obscured by poor test-retest reliability. However, as of yet, the psychometric properties of computational parameters have received limited attention (Ahn & Busemeyer, 2016; Browning et al., 2020; Nair et al., 2020; Paulus et al., 2016). Model-agnostic measures derived from reward processing tasks (e.g., percent correct) often have modest or poor reliability (Bland et al., 2016; Enkavi et al., 2019; Plichta et al., 2012). The few studies that have examined the reliability of PT and RL parameters based on gambling and various reward-processing tasks have reported either poor-to-good (Chung et al., 2017; Glockner & Pachur, 2012; Scheibehenne & Pachur, 2015) or poor reliability (Moutoussis et al., 2018; Shahar et al., 2019); although more recent studies have shown that higher reliability estimates can be achieved using hierarchical procedures (Brown et al 2020; Waltmann et al., 2022). Computational cognitive models are however often context-specific (Eckstein et al., 2022), suggesting that the reliability of computational decision-making processes may differ by task. However, no studies to date have reported the reliability of models derived from a bandit task with fluctuating (“restless”) reward and punishment probabilities (Daw et al., 2006) or an individually calibrated gambling task (Charpentier et al., 2017), despite these showing promise in computational psychiatry studies (e.g., Aylward et al., 2019; Charpentier et al., 2017).

A complementary perspective to understanding the reliability of computational cognitive models can be obtained through prediction. Generative models offer a substantial advantage in that they can both explain and predict behaviour. Unlike model-agnostic measures, computational parameters fit to one dataset should be able to predict future behaviour in the same individual. In other words, computational models can additionally be assessed by their ability to forecast future behaviour, equivalent to out-of-sample validation. This type of validation assesses model generalizability and is often referred to as predictive accuracy (Busemeyer & Wang, 2000; Glockner & Pachur, 2012; Scheibehenne & Pachur, 2015), but it has rarely been used as a metric of reliability. The aim of the current study was to assess the reliability of model-agnostic and computational parameters derived from two widely-used decision-making tasks (a restless four-armed bandit and a calibrated gambling task) using standard measures of stability and reliability (respectively, practice effects and intraclass correlations; ICCs) and additionally out-of-sample predictive accuracy for model parameters.

## Methods and Materials

### Participants

Fifty-four healthy participants were recruited from the UCL Institute of Cognitive Neuroscience Subject Database. Four participants were excluded for failing to complete the second session (final N=50: 32 females [64%]; age range=19-38; mean age=25.16, SD±5.48 years; mean education=17.38, SD=±3.24 years). Participants reported no current or past psychiatric or neurological disorder, cannabis use in the past 31 days, alcohol consumption in the past 24 hours, or any other recreational drug use in the week prior to participation. Participants provided written informed consent and were compensated at the end of their second session with a flat rate of £30 and a bonus of up to £20 based on task winnings. The study was approved by the UCL Psychology and Language Sciences Research Ethics Committee (Project ID Number: fMRI/2013/005) and was performed in accordance with the Declaration of Helsinki.

Sample size was determined by an *a priori* power analysis in G*Power (Faul et al., 2007). The power analysis was based on the smallest effect size of interest, r=0.4, since reliability below this threshold is conventionally considered poor (Fleiss, 2011). Detecting an effect size of this magnitude, at the one-tailed 0.05 alpha level with 90% power, requires 47 participants.

### Study procedure and tasks

Participants completed a battery of computerized tasks, including a restless four-armed bandit (Daw et al., 2006; Seymour et al., 2012) and an individually calibrated gambling task (Charpentier et al., 2017) over two sessions (mean test-retest interval = 13.96 days, SD=0.20). On each trial in the bandit task, participants chose one out of four bandits and received one out of four possible outcomes: reward, punishment, neither reward nor punishment or both reward and punishment (200 trials total). Win and loss probabilities fluctuated independently over time and between boxes (Figure 1a-b; Supplemental Materials). In the gambling task, participants chose between a 50-50 gamble and a sure option without receiving feedback. Trials were classified as either mixed (50% chance to win or lose money gamble, or a sure option of 0 points) or gain-only (50% chance to win or receive nothing gamble, or a variable sure gain; Figure 1c; Supplemental Materials). An initial training phase (50 mixed and 40 gain-only trials) was used to create individually calibrated offers (centred on indifference points) in a second phase (64 mixed and 56 gain-only trials). Calibration failed for one participant resulting in N=49 participants for this task. Both tasks lasted around 15 minutes.

**Figure 1.**
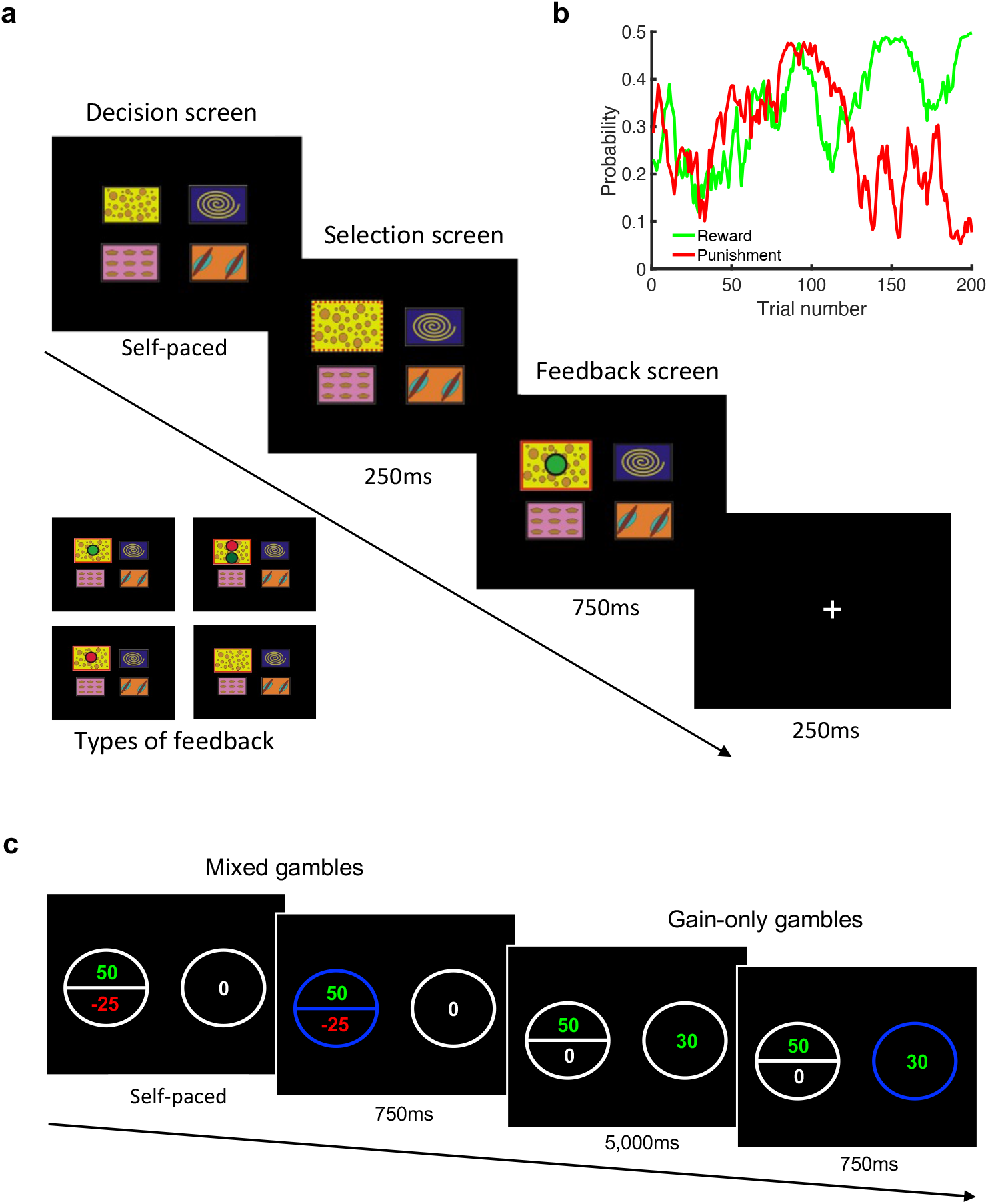
Four-armed bandit and gambling task. Example trial of the four-armed bandit task (a). On each trial, participants chose one out of four bandits and received one out of four possible outcomes: reward (green token), punishment (red token), neither reward nor punishment (empty box) or both reward and punishment (red and green token). An example of the win and loss probabilities fluctuating independently over time within one of the boxes (b). On each gambling task trial, participants chose between a 50-50 gamble and a sure (guaranteed amount of points) option (c). Trials were either mixed gambles (50-50 chance of winning or losing points or sure option of 0 points) or gain-only trials (50-50 chance of winning or receiving nothing or sure gain).

### Data analysis

Data were processed in Matlab (R2019b) and analysed in SPSS (v25, IBM Corp, Armonk, NY) and R (v. 4.2.1). Computational modelling was performed with the hBayesDM package for R (v.3.6.0; https://github.com/CCS-Lab/hBayesDM) (Ahn et al., 2017), which uses hierarchical Bayesian modelling in Stan (v.2.21.2). For all analyses, *p*<0.05 (two-tailed) was considered statistically significant. Cohen’s dz (within-subject) effect sizes are reported for practice effects (Lakens, 2013).

### Model-agnostic task analyses

Model-agnostic measures for the bandit task included the mean probability of repeating a choice after win-only, loss-only and no outcomes (‘neither’). A repeated-measures analysis of variance (ANOVA) was conducted with the within-subjects factors outcome (win, loss, neither) and session (session 1, session 2) to assess basic behaviour and practice effects.

Model-agnostic measures for the gambling task included the mean probability of gambling on mixed and gain-only trials. It was predicted that gambling would be higher on mixed trials, which was assessed using a repeated-measures ANOVA with within-subjects factors gamble (mixed, gain-only) and session (session 1, session 2).

As a supplementary analysis, bandit and gamble model-agnostic measures were additionally derived from trial-by-trial mixed-effects logistic regressions for each session since previous studies have suggested that hierarchically predicted values can improve the precision of reliability (Brown et al 2020; Waltmann et al., 2022). Further details on all model-agnostic measures are in Supplemental Materials.

### Computational modelling

The bandit task data was fit with seven different RL models previously described in detail (Aylward et al., 2019; Supplemental Materials). Three PT models were fit to the gambling task (Ahn et al., 2017; Charpentier et al., 2017; Kahneman & Tversky, 1979; Sokol-Hessner et al., 2009). Modelling was conducted on the second phase of the gambling task (i.e., on individually calibrated trials). The models were fit for each session separately, using separate hierarchical priors (group-level parameters), as this has shown to provide more accurate fits (Valton et al., 2020), and we wished to avoid artificially inflating reliability estimates. We also estimated model fits under a single hierarchical prior (session 1 and session 2 data together) as a sensitivity analysis (Supplemental Materials). Model comparison was performed with leave-one-out information criterion (LOOIC) where the winning model was the one with the lowest LOOIC. Several model validation checks were completed for the winning models, including examination of MCMC convergences, parameter recovery and recapitulation of real data (Daw, 2011; Kruschke, 2015; Wilson & Collins, 2019; Supplemental Materials; Figure S1-S8).

### Reliability analysis

Test-retest reliability was assessed with ICCs (ratios of intra-individual to inter-individual variability (Koo & Li, 2016; McGraw & Wong, 1996)), with values of <0.40 interpreted as poor, 0.4-0.6 as fair, 0.6-0.75 as good, and >0.75 as excellent reliability (Fleiss, 2011). A two-way mixed-effects model based on single-measures and absolute-agreement ICC was used (fixed effect: testing time-interval, random effect: subject), equivalent to ICC(A,1) (McGraw & Wong, 1996).

We also specified an additional model that considers data from both sessions jointly, taking the hierarchical structure into account and estimating reliability directly within the model, which has recently been shown to improve the estimation of reliability (Brown et al 2020; Haines et al., 2020; Waltmann et al., 2022). For model-agnostic measures, this was achieved by estimating a single mixed-effects logistic regression with random intercepts and slopes, accounting data for both sessions jointly by including session as a second-level grouping factor with subjects nested within session (Brown et al 2020; Waltmann et al., 2022). This allows extracting variance components from the logistic regression to calculate a one-way random-effects, absolute-agreement, single-measure ICC: ICC(1) in the McGraw and Wong (1996) convention. This was calculated for all bandit and gamble model-agnostic measures. For computational parameters, model-calculated Pearson’s r correlations were estimated between parameters from session 1 and 2 by fitting all the data from both sessions together and embedding a correlation matrix between sessions in the winning RL and PT models, using an identical approach described in detail previously (Haines et al., 2020; Pike et al., 2022). This essentially results in a multivariate prior where the correlation between group-level parameters across session are also considered and the uncertainty of point estimates are accounted for in the model-calculated reliability estimate. Separate subject-specific parameters are estimated for each session as well, but these were not extracted to calculate reliability on as previous studies have suggested that this overestimates reliability (Waltmann et al., 2022).

### Posterior predictive performance

To assess to what extent an individual’s future behaviour can be predicted using a generative model fit to their own task performance two weeks earlier, we calculated the probability of participants’ choices on each trial (i.e., the softmax output), given their session 2 data and model parameter estimates from session 1. Probabilities were averaged across trials for each individual.

Since hierarchical parameter estimation produces ‘shrinkage’, effectively pulling parameter estimates from different individuals closer to each other (which improves estimation accuracy), it is possible that future performance may also be predicted above-chance using other participants’ parameter estimates from session 1 (e.g., participant A’s parameter estimates from session 1 predicting participant B’s session 2 choices). We therefore assessed whether using an individual’s model parameter estimates from session 1 predicted the same individual’s choices on session 2 better than using all other subjects’ model parameter estimates. To construct the latter measure, for each subject, we predicted trial-by-trial choices on session 2 based on parameter estimates from every other participant’s session 1 model, and averaged the probabilities across all participants. Additionally, we compared subjects’ own session 1 parameters in predicting their session 2 behaviour to the mean session 1 model parameter priors in predicting future behaviour.

### Data accessibility

All script code and data are available on OSF at https://osf.io/n7czx/.

## Results

### Four-armed bandit task: model-agnostic results

#### Basic behaviour and practice effects

As expected, there was a main effect of outcome type on behaviour (*F*_(2,98)_=117.39, *p*<0.001, 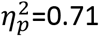; Figure 2a). The probability to repeat a choice was significantly greater after wins compared with both losses and outcomes on which neither wins nor losses occurred, and greater after neither compared with losses (all *p*<0.001). There was no significant main effect of testing session (*F*_(1, 49)_=0.01, *p*=0.91, 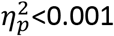). There was a significant outcome-by-session interaction (*F*_(2,98)_=3.12, *p*=0.049, 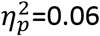), reflecting slightly increased repeated choices after wins and decreased repeated choices after losses. However, the difference in the tendency to repeat a choice between session 1 and session 2 did not reach significance following any of the outcome types (loss: t_(49)_=1.45, *p*=0.15, d_z_=0.21; win: t_(49)_=0.87, *p*=0.39, d_z_=0.12; neither: t_(49)_=0.54, *p*=0.59, d_z_=0.08), and therefore we do not interpret this result further.

**Figure 2:**
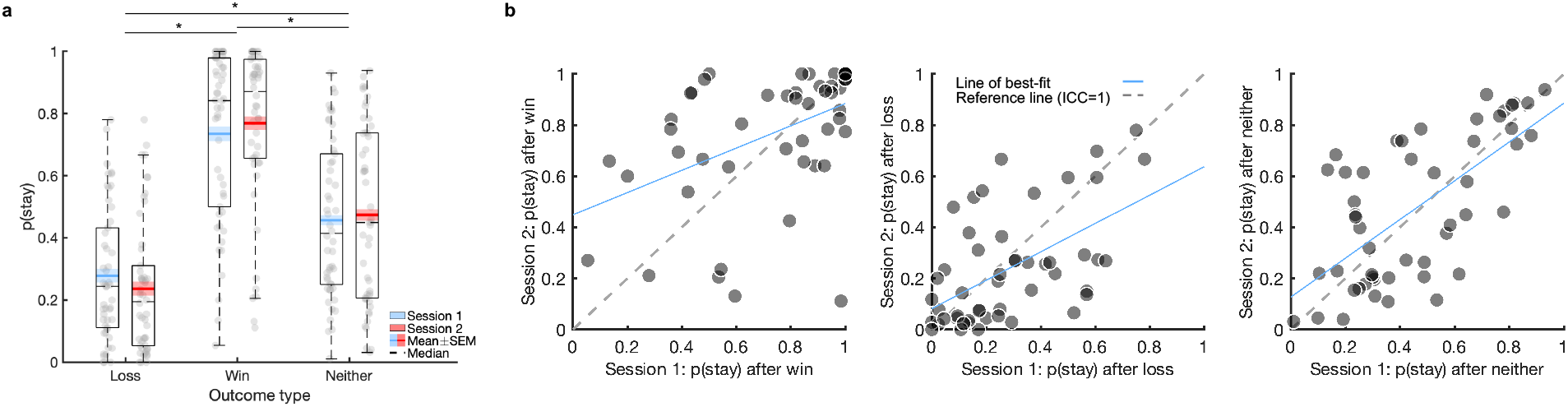
Basic behaviour, practice effects, and test-retest reliability of model-agnostic measures on the four-armed bandit task. Boxplots of the four-armed bandit task showing probability to stay after a certain outcome in session 1 and 2 (a). The probability to stay was significantly different after each outcome type (Loss<Neither<Win) but no clear practice effect was evident. Scatter plots of the model-agnostic measures comparing behaviour on two testing sessions approximately 2 weeks apart (b). Lightly shaded regions in Figure 2a represent within-subjects standard error of the mean (SEM). \**p*<0.001

#### Test-retest reliability

The model-agnostic measures exhibited fair-to-good reliability (Figure 2b; Table 1), which did not improve substantially when examined in separate trial-by-trial mixed logistic regressions (Table S1). However, reliability did increase, particularly for p(stay) after win and loss when estimated as part of a joint mixed-effects logistic regression (Table 1).

**Table 1:**
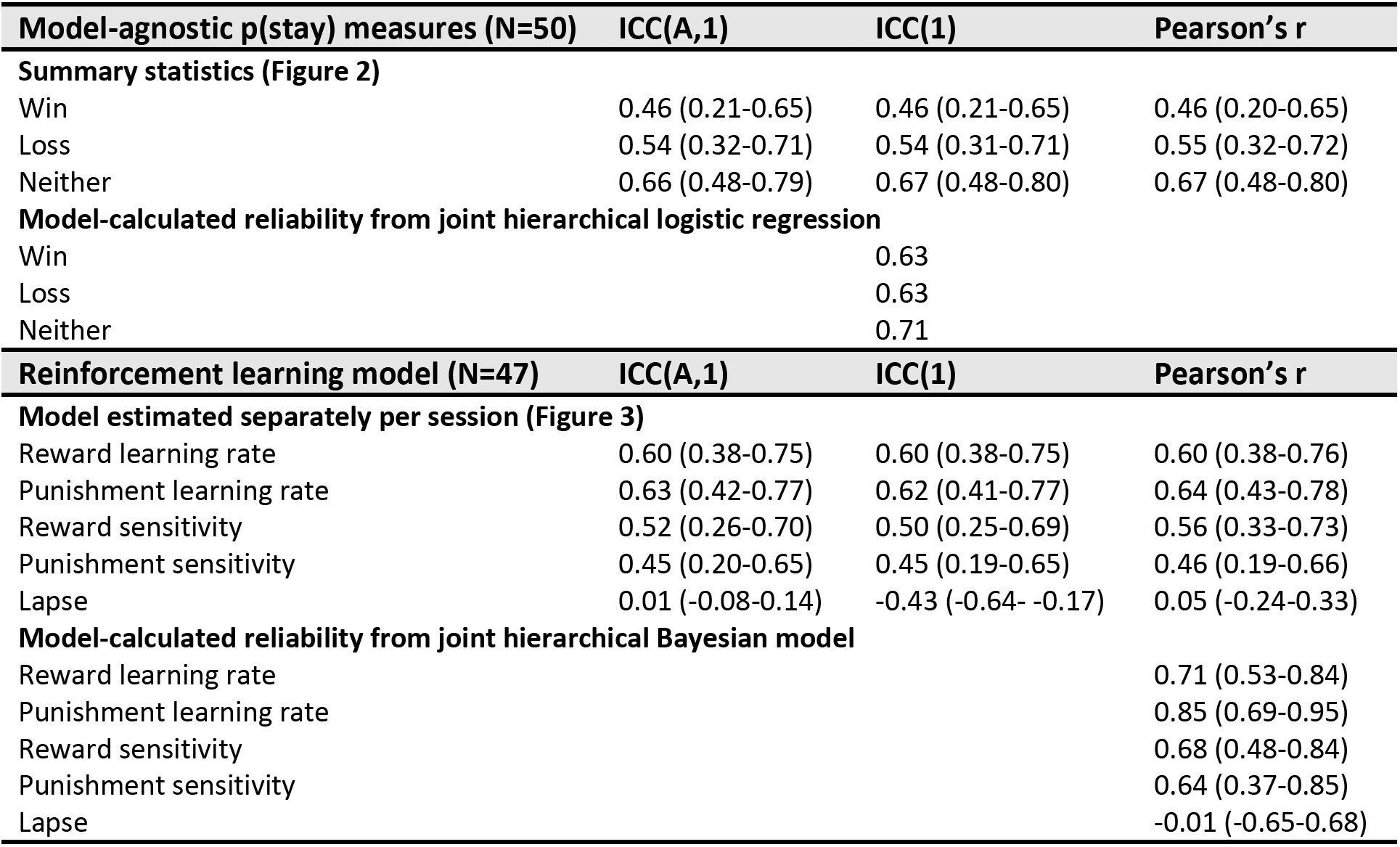
Reliability of model-agnostic and computational measures of the four-armed bandit task. All measures but the lapse parameter are significant at *p*<0.05. Brackets represent the 95% confidence interval.

| Model-agnostic p(stay) measures (N=50) | ICC(A,1) | ICC(1) | Pearson's r |
| --- | --- | --- | --- |
| **Summary statistics (Figure 2)** |  |  |  |
| Win | 0.46 (0.21-0.65) | 0.46 (0.21-0.65) | 0.46 (0.20-0.65) |
| Loss | 0.54 (0.32-0.71) | 0.54 (0.31-0.71) | 0.55 (0.32-0.72) |
| Neither | 0.66 (0.48-0.79) | 0.67 (0.48-0.80) | 0.67 (0.48-0.80) |
| **Model-calculated reliability from joint hierarchical logistic regression** |  |  |  |
| Win |  | 0.63 |  |
| Loss |  | 0.63 |  |
| Neither |  | 0.71 |  |
| Reinforcement learning model (N=47) | ICC(A,1) | ICC(1) | Pearson's r |
| **Model estimated separately per session (Figure 3)** |  |  |  |
| Reward learning rate | 0.60 (0.38-0.75) | 0.60 (0.38-0.75) | 0.60 (0.38-0.76) |
| Punishment learning rate | 0.63 (0.42-0.77) | 0.62 (0.41-0.77) | 0.64 (0.43-0.78) |
| Reward sensitivity | 0.52 (0.26-0.70) | 0.50 (0.25-0.69) | 0.56 (0.33-0.73) |
| Punishment sensitivity | 0.45 (0.20-0.65) | 0.45 (0.19-0.65) | 0.46 (0.19-0.66) |
| Lapse | 0.01 (-0.08-0.14) | -0.43 (-0.64- -0.17) | 0.05 (-0.24-0.33) |
| **Model-calculated reliability from joint hierarchical Bayesian model** |  |  |  |
| Reward learning rate |  |  | 0.71 (0.53-0.84) |
| Punishment learning rate |  |  | 0.85 (0.69-0.95) |
| Reward sensitivity |  |  | 0.68 (0.48-0.84) |
| Punishment sensitivity |  |  | 0.64 (0.37-0.85) |
| Lapse |  |  | -0.01 (-0.65-0.68) |

### Four-armed bandit task: modelling results

Model comparison indicated that the winning (most parsimonious) model was the five-parameter ‘Bandit4arm_lapse’ (nomenclature from the hBayesDM package) model, with reward and punishment learning rate parameters, reward and punishment sensitivity parameters and a lapse parameter (final parameter captures random responding; Table S2), consistent with previous reports (Aylward et al., 2019). Three individuals were excluded due to difficulties in obtaining mean parameter estimates, as multiple peaks were evident in the posterior distribution of at least one parameter (Figure S2). The Bandit4arm_lapse model was therefore re-fit without these participants. Excluding these participants did not affect test-retest reliability inference.

#### Practice effects

On session 2 there were significant increases in the reward sensitivity (t_(46)_=3.00, *p*=0.004, d_z_=0.44) and lapse parameters (t_(46)_=8.88, *p*<0.001, d_z_=1.29), but not on any of the other parameters (reward learning rate: t_(46)_=1.28, *p*=0.21, d_z_=0.19; punishment learning rate: t_(46)_=1.74, *p*=0.09, d_z_=0.25; punishment sensitivity: t_(46)_=1.28, *p*=0.21, d_z_=0.19; Figure 3a). However, there were no significant practice effects when the data was fit under a single hierarchical prior (Table S3).

**Figure 3:**
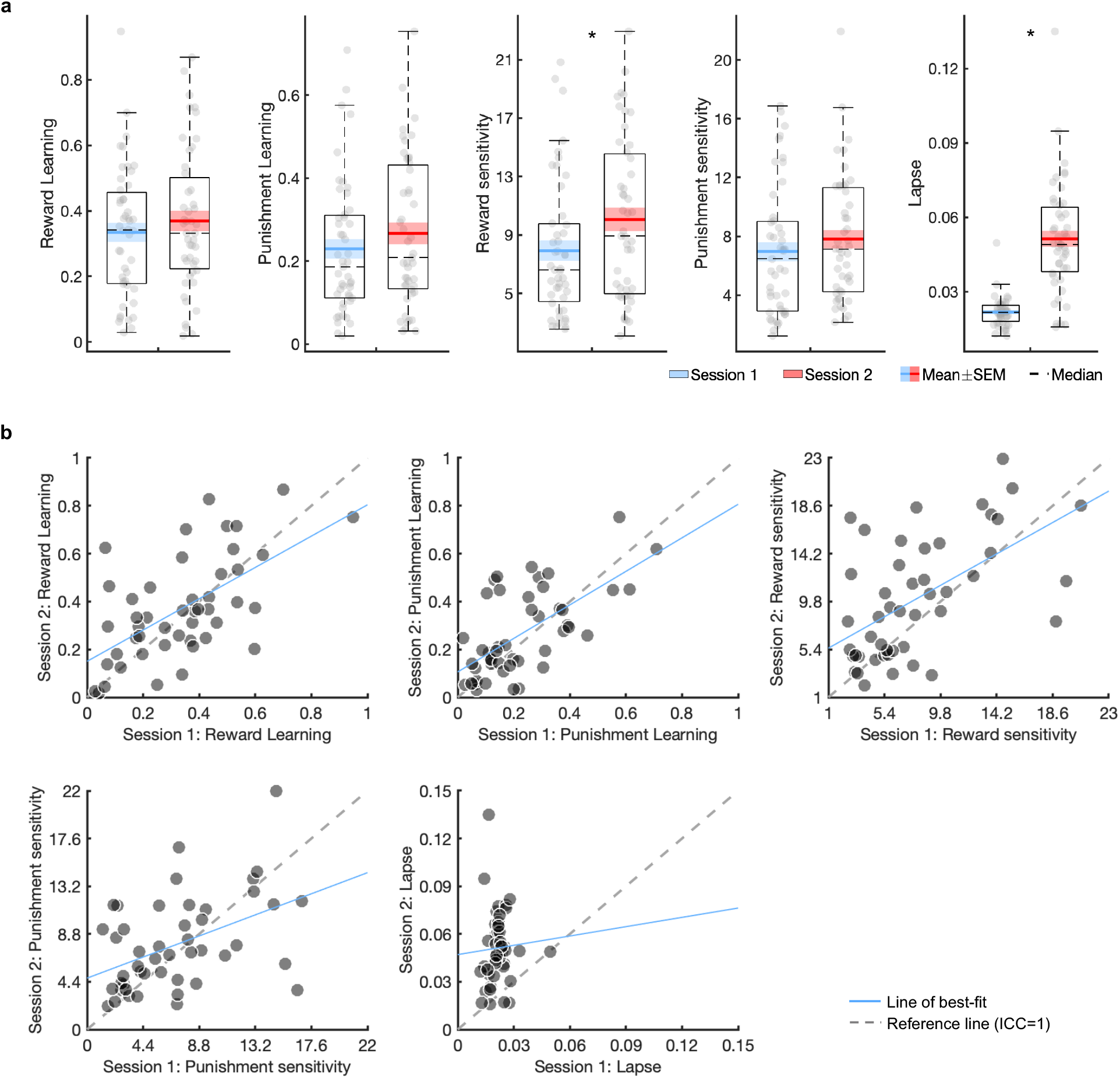
Practice effects and test-retest reliability of the winning reinforcement learning model parameters derived from the four-armed bandit task. Boxplots show point estimates of the Bandit4arm_lapse model parameters in session 1 and 2, fit under separate priors (a). Scatter plots of the Bandit4arm_lapse model parameters over session 1 and 2 are presented (b). SEM: standard error of the mean. \**p*<0.05.

#### Test-retest reliability

All estimated Bandit4arm_lapse model parameters, except the lapse parameter, demonstrated fair-to-good reliability (Figure 3b; Table 1). This did not substantially change when parameters were estimated under a single hierarchical prior (Table S3). However, examining the correlation between parameters as estimated within a generative joint model showed good-to-excellent reliability, improving reliability across all but the lapse parameter (Table 1).

#### Posterior predictive performance

Parameter estimates from session 1 predicted task performance on session 2 substantially better than chance (mean = 42%, chance = 25% accuracy; t_(46)_=9.10, *p*<0.001; Figure 4a), indicating that the model could predict future choices by using a generative model fit to the same participants’ data two weeks earlier. Using an individual’s parameter estimates to predict their own future choices was significantly better than when that prediction was based on the average of the other participants’ session 1 estimates (t_(46)_=3.20, *p*=0.003; Figure 4b). However, there was no significant difference between using an individual’s own session 1 parameter estimates compared with the session 1 mean prior parameter in predicting future behaviour (t_(46)_=1.04, *p*=0.30; Figure 4c).

**Figure 4.**
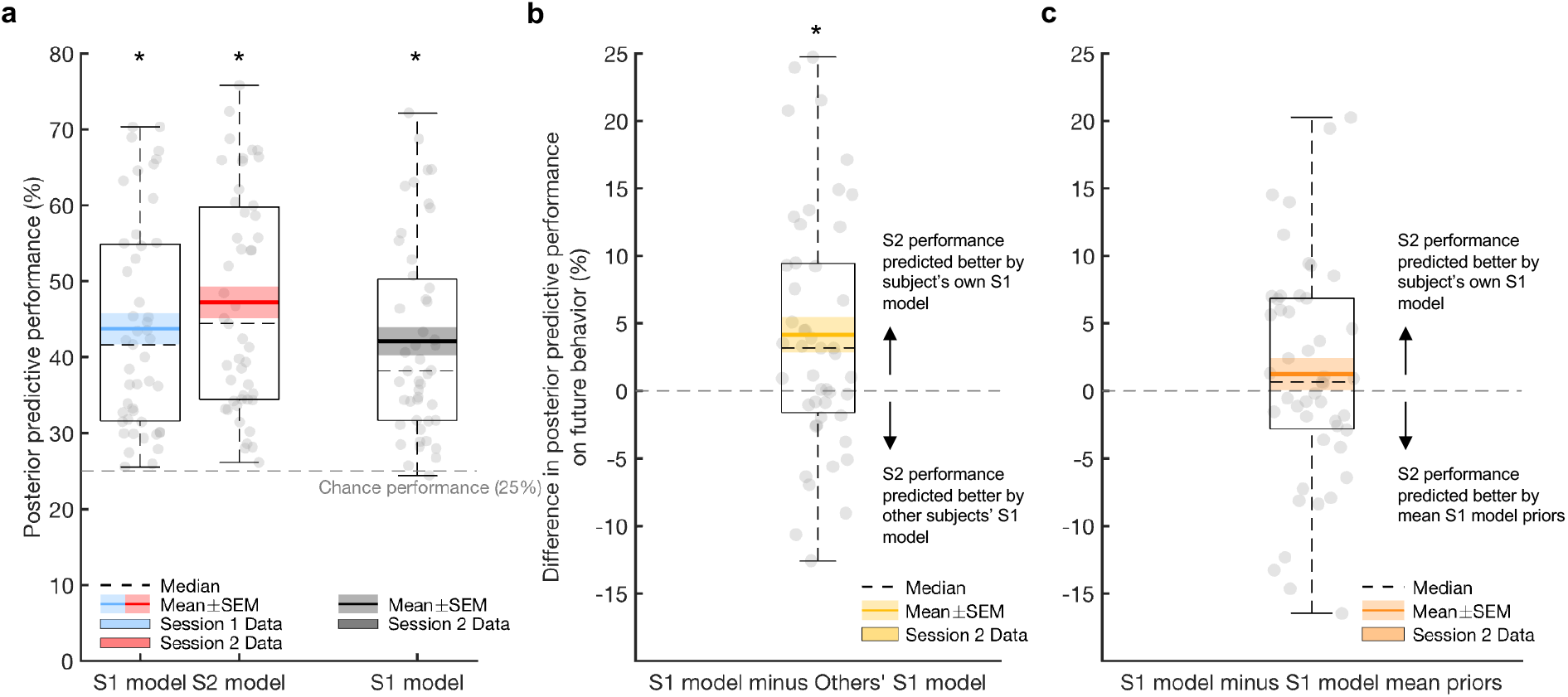
Posterior predictive performance of the winning reinforcement learning model derived from the four-armed bandit task. Boxplots depicting accuracy of bandit4arm_lapse model in predicting choices (a). Model estimates from session 1 (S1) predicted future session 2 (S2) behaviour above chance (black boxplot). Both S1 and S2 model estimates also predicted behaviour on the same session significantly above chance (blue and red boxplots). Predicting future performance (session 2 data) using a participant’s own model parameter estimates was significantly better than using other participants’ S1 model parameter estimates (b) but not when comparing against the mean S1 model priors (c). SEM: standard error of the mean. \**p*<0.01.

### Gambling task: model-agnostic results

#### Basic behaviour and practice effects

As expected, propensity to gamble was significantly higher on mixed gambles (*F*_(1, 48)_=13.71, *p*=0.001, 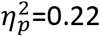). There were no significant main (*F*_(1, 48)_=0.76, *p*=0.40, 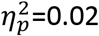) or interaction (*F*_(1, 48)_=1.07, *p*=0.31, 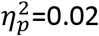) effects of session on the propensity to gamble (session differences: probability to gamble on mixed trials t_(48)_=0.23, *p*=0.82, d_z_=0.03; probability to gamble on gain-only trials t_(48)_=1.51, *p*=0.14, d_z_=0.22; Figure 5a).

**Figure 5:**
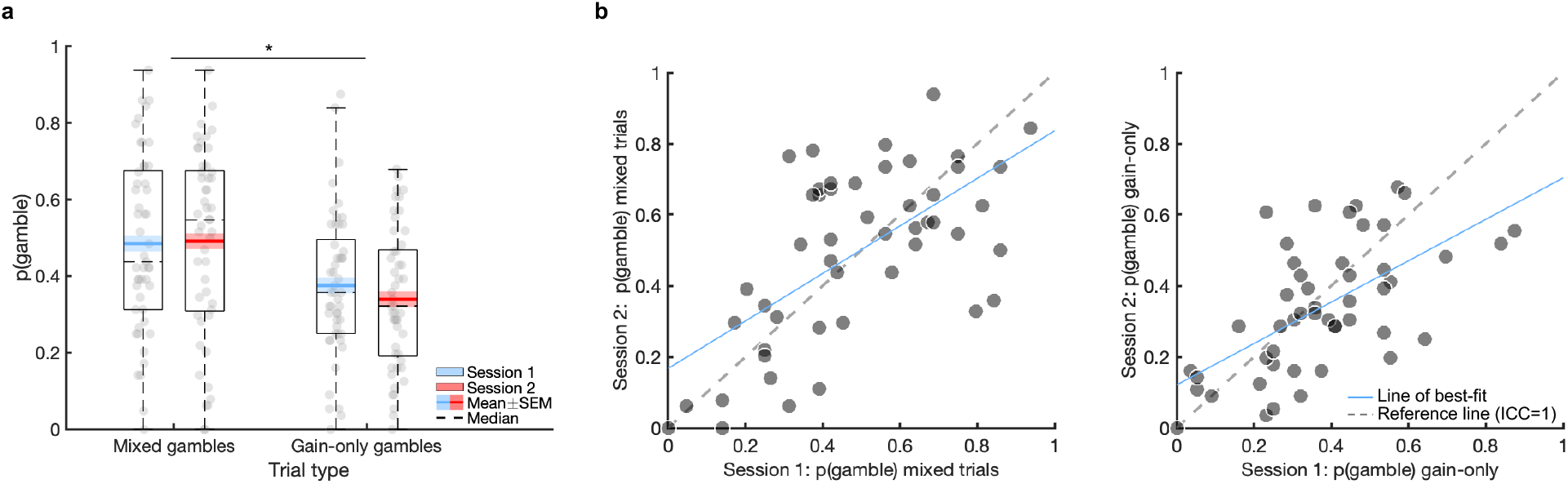
Basic behaviour, practice effects, and test-retest reliability of model-agnostic measures on the gambling task. Boxplots show the probability to gamble based on the trial type in session 1 and 2, with no significant session effects (a). Scatter plots of the model-agnostic measures over session 1 and 2 (b). Lightly shaded regions in Figure 5a represent within-subjects standard error of the mean (SEM). \**p*<0.001.

#### Test-retest reliability

Model-agnostic measures on the gambling task exhibited fair-to-good reliability (Figure 5b; Table 2) and did not change substantially when examined hierarchically in separate mixed logistic regressions (Table S4). Calculating reliability within a joint mixed logistic regression numerically improved the reliability of the gambling model-agnostic measures (Table 2).

**Table 2:**
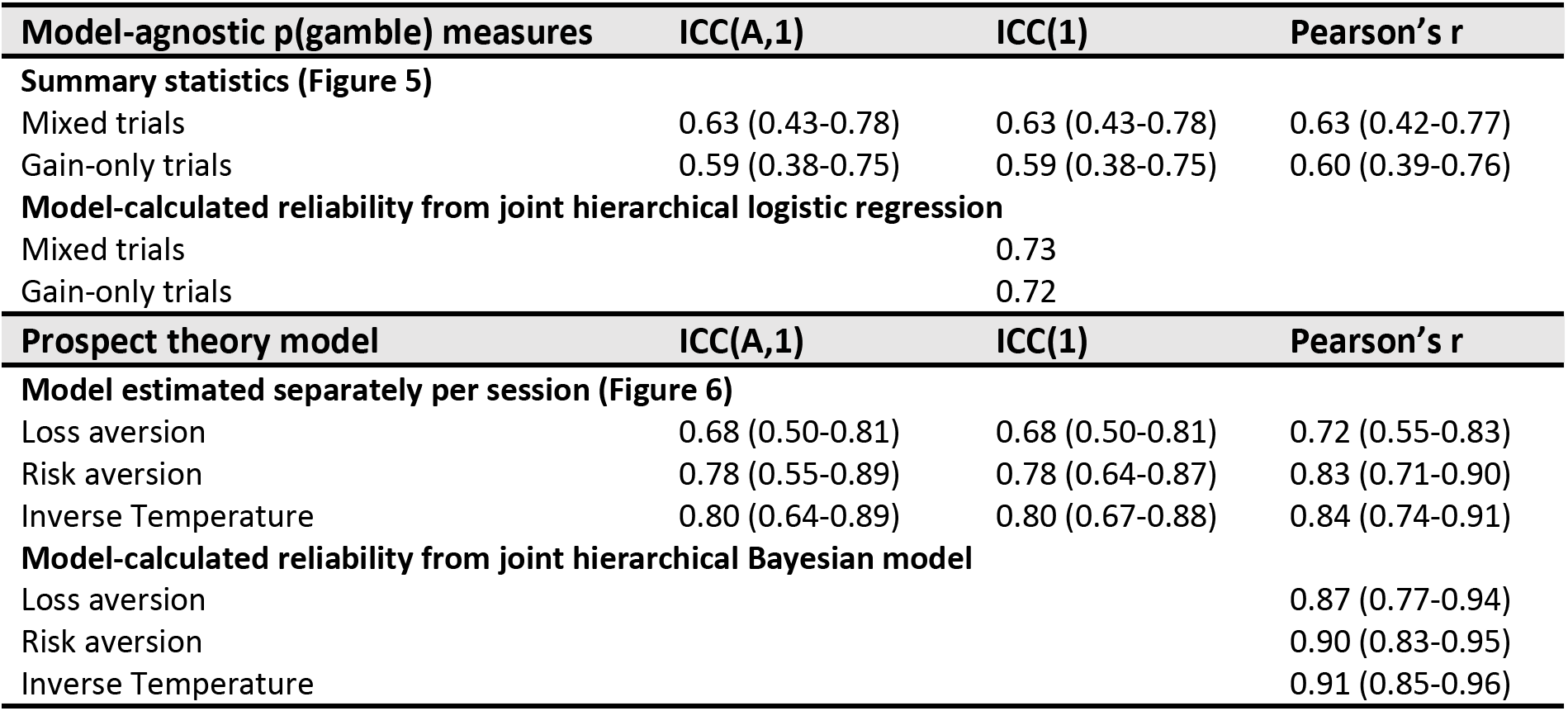
Reliability of model-agnostic and computational measures of the gambling task. All measures are significant at *p*<0.05. Brackets represent the 95% confidence interval.

| Model-agnostic p(gamble) measures | ICC(A,1) | ICC(1) | Pearson's r |
| --- | --- | --- | --- |
| **Summary statistics (Figure 5)** |  |  |  |
| Mixed trials | 0.63 (0.43-0.78) | 0.63 (0.43-0.78) | 0.63 (0.42-0.77) |
| Gain-only trials | 0.59 (0.38-0.75) | 0.59 (0.38-0.75) | 0.60 (0.39-0.76) |
| **Model-calculated reliability from joint hierarchical logistic regression** |  |  |  |
| Mixed trials |  | 0.73 |  |
| Gain-only trials |  | 0.72 |  |
| Prospect theory model | ICC(A,1) | ICC(1) | Pearson's r |
| **Model estimated separately per session (Figure 6)** |  |  |  |
| Loss aversion | 0.68 (0.50-0.81) | 0.68 (0.50-0.81) | 0.72 (0.55-0.83) |
| Risk aversion | 0.78 (0.55-0.89) | 0.78 (0.64-0.87) | 0.83 (0.71-0.90) |
| Inverse Temperature | 0.80 (0.64-0.89) | 0.80 (0.67-0.88) | 0.84 (0.74-0.91) |
| **Model-calculated reliability from joint hierarchical Bayesian model** |  |  |  |
| Loss aversion |  |  | 0.87 (0.77-0.94) |
| Risk aversion |  |  | 0.90 (0.83-0.95) |
| Inverse Temperature |  |  | 0.91 (0.85-0.96) |

### Gambling task: modelling results

The winning model was the PT model (‘ra_prospect’ in the hBayesDM package) with loss aversion, risk aversion and inverse temperature parameters (last parameter represents choice consistency; Table S5), consistent with previous reports (Charpentier et al., 2017). A loss aversion parameter above 1 represents overweighting of losses to gains, while a risk aversion parameter less than 1 indicates aversion to risk. Neither test-retest nor practice effects were substantially altered when the model was fit under a single hierarchical prior (Table S6).

#### Practice effects

There were significant session effects on all PT model parameters (on session 2: decreased loss aversion: t_(48)_=2.17, *p*=0.04, d_z_=0.31; decreased risk aversion: t_(48)_=4.04, *p*<0.001, d_z_=0.58; increased inverse temperature: t_(48)_=3.07, *p*=0.004, d_z_=0.44; Figure 6a).

**Figure 6:**
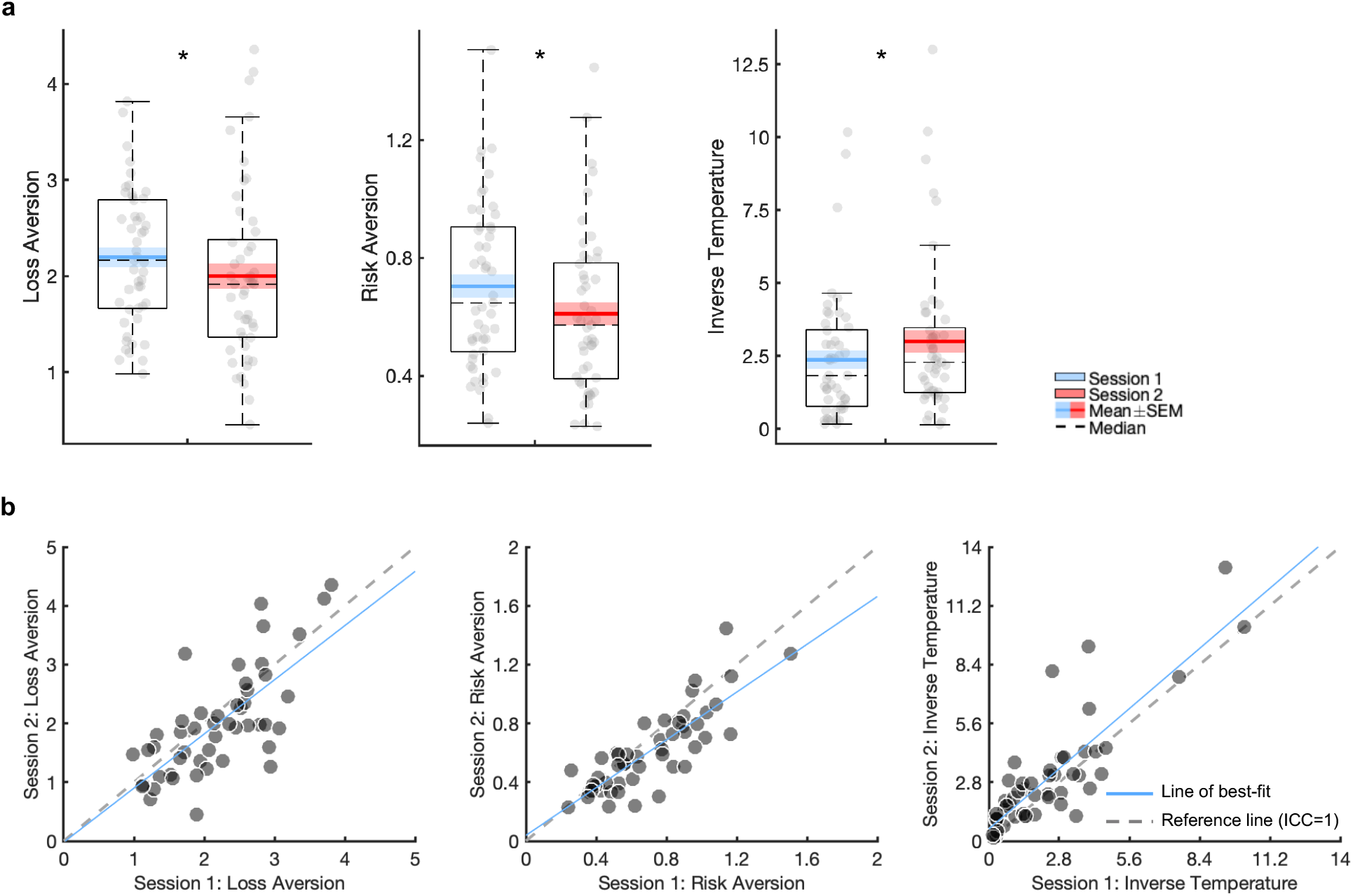
Practice effects and test-retest reliability of the prospect theory model derived from the gambling task. Boxplots show point estimates of the prospect theory model parameters in session 1 and 2, fit under separate priors (a). Scatter plots of the prospect theory model parameters over session 1 and 2 are presented (b). SEM: standard error of the mean. \**p*<0.05.

#### Test-retest reliability

All estimated parameters demonstrated good-to-excellent reliability (Figure 6b; Table 2), and showed excellent reliability when estimating a correlation matrix within a joint model (Table 2).

#### Posterior predictive performance

PT model parameters from session 1 predicted future choices at session 2 significantly above chance (mean = 68%, chance = 50% accuracy; t_(48)_= 12.08, *p*<0.001; Figure 7a). Predicting future performance at session 2 was significantly higher when based on participants’ own parameter estimates from session 1 compared with model parameter estimates of other participants from session 1 (t_(48)_=8.38, *p*<0.001; Figure 7b). This was also true when comparing between participants’ own session 1 parameter estimates with the mean session 1 prior for parameters (t_(48)_=6.28, *p*<0.001; Figure 7c).

**Figure 7:**
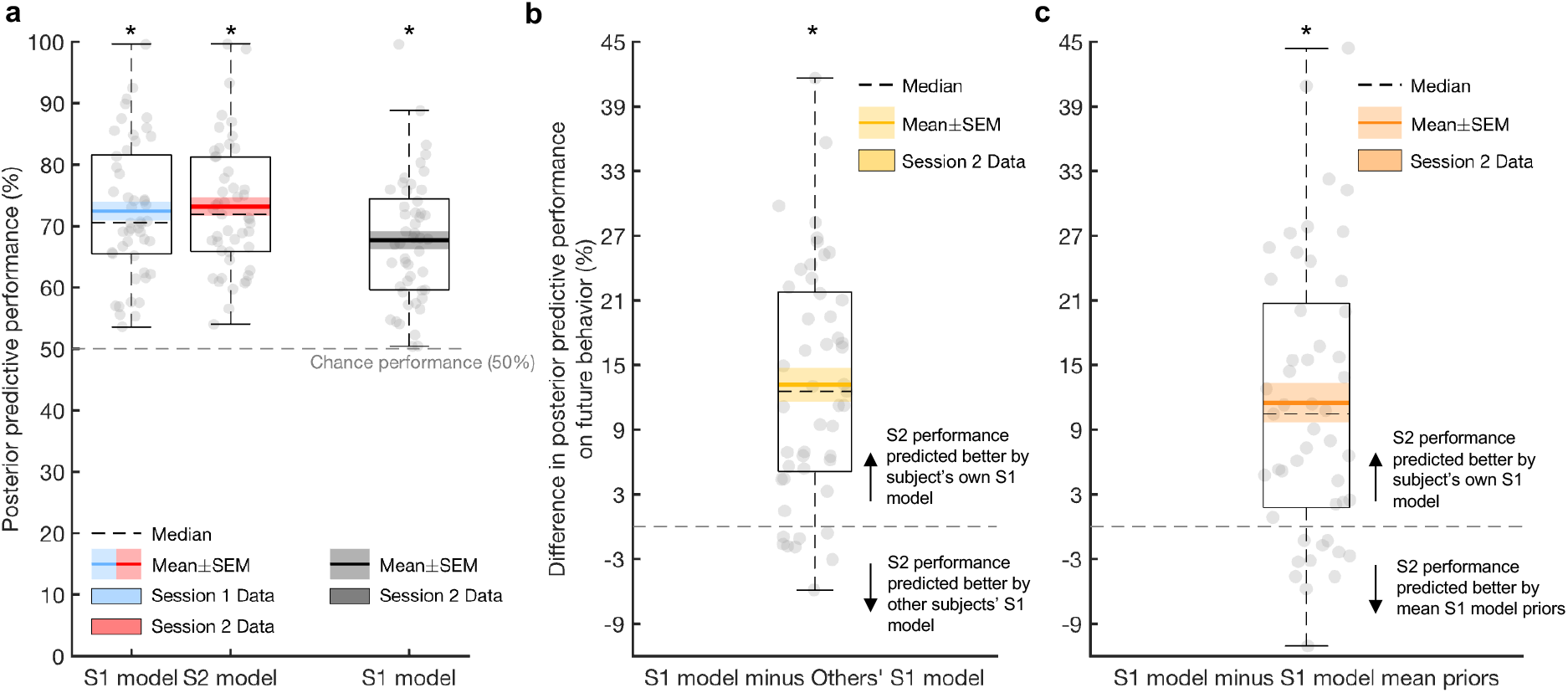
Posterior predictive performance of the prospect theory model derived from the gambling task. Boxplots depicting accuracy of prospect theory model in predicting choices (a). Session 1 (S1) model estimates predicted S1 behaviour significantly above chance (blue boxplot), as did session 2 (S2) model estimates on S2 data (red boxplot). Importantly, model parameter estimates from S1 predicted task performance from S2 above chance (black boxplot). Predicting future S2 performance using a participant’s own S1 model parameter estimates was significantly better than using other participants’ S1 model parameter estimates (b) and mean S1 model priors (c). SEM: standard error of the mean. \**p*<0.001.

## Discussion

Reliability has garnered increased attention in recent years, with worryingly low reliability across conventional measures from cognitive tasks and functional neuroimaging (Elliott et al., 2020; Enkavi et al., 2019; Noble et al., 2019; Nord et al., 2017; Rodebaugh et al., 2016). However, fewer attempts have been made to examine the reliability of computational cognitive measures. Here we assessed the psychometric properties of computational models derived from a restless four-armed bandit and a calibrated gambling task. Overall, most parameters reflecting RL and decision-making processes exhibited adequate reliability and predicted future performance well. These results provide promise for their use in clinical settings. However, this conclusion depends on the specific parameters assessed in each task, highlighting the complexities of translating tasks to the clinic.

### Four-armed bandit RL model reliability

Reward and punishment learning rates from the bandit task demonstrated good reliability while reward and punishment sensitivity showed fair reliability, suggesting that this task may be more suitable for assessing learning rates than sensitivity. Elevated punishment learning rates (faster learning in the face of negative outcomes) and lapse values have been associated with greater mood and anxiety symptoms, representing potential measurable mechanistic treatment targets (Aylward et al., 2019). However, the present study suggests that the lapse parameter, which exhibited poor reliability as assessed by the bandit task, may not be a suitable target. This parameter measures responding not captured by the model (including goal-directed and random exploration), and the sources of this ‘noise’ might differ across sessions. It is therefore perhaps unsurprising that this parameter was unreliable. Crucially, the lapse parameter showed poor recoverability, which places an upper limit on its potential reliability. Some of this poor recoverability may be explained by limited lapse variation, especially in session 1 (Figure S3). The distribution of the group-level standard deviation lapse parameter was biased towards smaller values here such that the lapse parameter did not vary greatly between individuals (Figure S5). This suggests that the lapse parameter could be replaced with a constant and inference on this parameter is not advised.

Although no prior studies have specifically investigated ICC properties of the current RL model, one previous study found similarly poor reliability of the lapse parameter across six months in a go/no-go RL model in adolescents (Moutoussis et al., 2018). In contrast to our results, this study also reported poor reliability of both reward and punishment learning rates. These differences may arise for a multitude of reason, such as using different tasks (an orthogonalised go/no-go task versus a restless bandit task), testing time-windows (six months versus two weeks), populations (adolescents versus adults), or models. It is not possible to delineate these diverging results without systematically comparing these factors in one study. Importantly, however, we provide evidence that it is possible to achieve at least moderate reliability for some canonical RL parameters.

Interestingly, the model-agnostic outcome measures of the bandit task exhibited similar reliability to the computational measures. Model-agnostic measures of cognitive tasks have often been reported to exhibit poor-to-moderate reliability (Enkavi et al., 2019; Hedge et al., 2018; Plichta et al., 2012; Rodebaugh et al., 2016). It has been argued that this may be due to their inability to capture the generative process underlying task performance (Huys et al., 2021; Price et al., 2019). Our results suggest that it should not be assumed that computational parameters will always provide greater reliability than non-computational ones. However, the model-agnostic outcome measures are only a proxy of the processes the bandit task assesses, as it is difficult to compute model-agnostic equivalents of some parameters, such as reward/punishment sensitivity. Indeed, models make explicit and falsifiable predictions of the components driving behaviour, which can be refined and used to simulate artificial data to generate new predictions. Thus, computational modelling is a more rigorous and preferable method for assessing behaviour than model-agnostic measures, which unlike computational methods, lack the mechanistic insights into the underlying processes generating behaviour.

### Gambling PT model reliability

While reliability of parameters ranged from poor-to-good in the bandit task, parameters from the gambling task showed good-to-excellent test-retest reliability. These were also higher than the reliability of the model-agnostic measures, suggesting that computational models may offer advantages in psychometric properties here. In particular, the risk aversion parameter, which has previously been associated with anxiety (Charpentier et al., 2017), exhibited excellent reliability (ICC=0.78), providing promise for use in clinical research. These results show higher reliability than previous studies (loss aversion r≈0.25-0.61, risk aversion r≈0.50-0.60, inverse temperature r≈0.30-0.60; Chung et al., 2017; Glockner & Pachur, 2012; Scheibehenne & Pachur, 2015). These studies all used different estimation procedures, including hierarchical Bayesian, and employed both longer and shorter testing time-windows than the current study, suggesting that these factors may not fully explain the differences. It is possible that our results instead stem from different PT model specifications, as well as different task designs. Indeed, a strength of the gambling task is that we calibrated offers to each individual’s indifference point (Charpentier et al., 2017). A similar approach of dynamically updating parameter values to each individual during task performance has previously been suggested as a solution to unreliable cognitive tasks (Palminteri & Chevallier, 2018). This may allow for removing any potential state influences across participants to extract more trait-like measures of the variables of interest (here risk/loss aversion).

### Predictive accuracy

We also examined how well the models predicted future task performance, which provides complementary perspective on reliability, unique to computationally-informed measures. Notably, for the PT model participants’ own parameter estimates from the first session were better at predicting their future performance compared with using parameter estimates from all other participants and from model priors. Individuals’ own RL parameters only provided an advantage in predicting future performance when compared with other’s parameter estimates but not model priors. It is likely that the RL model did not perform as well on this metric as the PT model because the RL model does not provide as close a fit to behaviour, potentially due to some participants performing at chance level. Overall, this indicates that individuals may also reliably differ in the cognitive mechanisms underlying their decisions, and offers reassurance that hierarchical estimation procedures are suitable for inter-individual inferences (Brown et al., 2020; Daw, 2011; Scheibehenne & Pachur, 2015). In other words, individuals show relatively unique computational decision-making profiles, particularly in the PT model. This is consistent with two previous studies using a different PT model and gambling task (Glockner & Pachur, 2012; Scheibehenne & Pachur, 2015).

### Implications

The RL parameters showed relatively modest reliability, suggesting that these processes are more vulnerable to state influences or measured with less precision than PT parameters. A consequence of this is that larger sample sizes may be required to examine effects, as effect sizes would be expected to be lower, relative to PT tasks (Fleiss, 2011). Interpreting the marked difference in reliability between the PT and RL models is not straightforward, as these models measure complementary aspects of cognition. The bandit task is a learning paradigm that requires constant updating of optimal choices. It is possible that in the first session individuals had not yet stabilized on a cognitive strategy and were still learning the task structure, as indicated by lower evidence of the winning model in session 1 compared with session 2 (Table S2). It would be interesting to explore if an initial baseline session, would improve test-retest reliability.

It should also be noted that we observed substantially greater reliability of RL (good-to-excellent) and PT (excellent) parameters when estimated within the joint generative model, in line with previous studies (Brown et al., 2020; Waltmann et al., 2022). Similarly, calculating the reliability of model-agnostic measures directly from a joint logistic regression improved model-agnostic reliability estimates, although in the range of good reliability. Similar to the hierarchical estimation approach of separate sessions, joint estimation regularises estimates toward the group mean for that session, improving the accuracy of point estimates. However, joint estimation additionally considers the potentially correlated structure. While this can increase the within-subject precision, it can consequently inflate the reliability of point estimates extracted from joint models (hence why reliability was not calculated on the point estimates in these models; Waltmann et al., 2022). Estimating reliability directly within the model, however, takes the uncertainty around the point estimates into account, providing unbiased reliability estimates (Waltmann et al., 2022). This suggests that the increased precision (and here thus improved reliability) may stem from the greater information (including more data points) available in joint modelling. This analysis approach may therefore be preferred when the study design allows for it.

### Limitations

A potential limitation of our study is the sample tested, as the reliability of tasks in healthy individuals may differ from that in clinical groups. Similarly, our results only speak to reliability over two weeks. Thus, it is possible that longer time periods may produce lower reliability, which should be assessed in future studies. Reliability over two weeks is particularly informative for interventional studies such as randomised controlled trials (e.g., for rapid-acting antidepressants or for early markers of response for traditional antidepressants/ psychotherapies). This time-window is also in line with other reliability studies aiming to establish reliability of measures for e.g., individual differences (e.g., Hedge et al., 2018; Nord et al., 2017), based on the assumption that measures should remain relatively stable over a short time-period.

## Conclusion

In summary, we show that commonly-used computational parameters derived from an RL ‘restless’ bandit task and a calibrated gambling task exhibit fair-to-excellent reliability. Specifically, learning rates showed good reliability and sensitivity parameters showed fair reliability from the RL model, while loss aversion had good reliability and risk aversion and inverse temperature displayed excellent reliability from the PT model. These models can further be used to predict future behaviour in the same individuals, especially PT model parameters, indicating that the decision-making processes assessed in these tasks represent relatively consistent and unique characteristics of an individual. These findings take us one step closer to translating computational measures of behaviour into clinical application.

## Acknowledgements

This work was supported by a Wellcome Trust - NIH PhD studentship (200934/Z/16/Z) to A.M.; a National Institute for Health Research (NIHR) Biomedical Research Center (BRC) fellowship to V.V.; and by a Wellcome Investigator Award (101798/Z/13/Z) to J.P.R. The funders had no role in the study design, data collection and analysis, decision to publish or preparation of the manuscript. For the purpose of Open Access, the author has applied a CC BY public copyright licence to any Author Accepted Manuscript version arising from this submission. The authors thank Eoin Travers and Oliver J. Robinson for valuable input on an earlier version of the manuscript. A preprint was previously published at bioRxiv (https://doi.org/10.1101/2021.06.30.450026).

## Authors’ contribution

A.M. and J.P.R. conceived and designed the study. A.M. acquired the data and performed analyses under the supervision of V.V. and J.P.R. All authors contributed to interpreting the data and drafting or substantially revising of the manuscript.

