## Supplemental Materials for "Reliability of Decision-Making and Reinforcement Learning Computational Parameters" (from another version of this preprint: version 3, 2022-09-02)

### Supplementary Materials

#### Study procedure

Tasks were presented on a laptop using MATLAB (R2015b, The MathWorks, Inc., Natick, MA, United States) with Cogent (Wellcome Trust Centre for Neuroimaging and Institute of Cognitive Neuroscience, UCL, London, U.K.). Participants completed a battery of eight tasks, including a restless four-armed bandit and a calibrated gambling task. Tasks were presented in a random order in each session, for each participant. Participants were informed that at the end of each session the computer would randomly pick 100 trials across all the tasks to calculate the bonus.

#### General computational modelling methods

Model fitting was achieved through hierarchical Bayesian modelling using Markov chain Monte Carlo (MCMC) sampling to estimate the posterior distribution of model parameter values. Hierarchical model-fitting uses group-level information to inform individual parameter estimates and has shown to provide more accurate parameter estimates than non-hierarchical Bayesian model-fitting procedures (Ahn et al., 2011; Brown et al., 2020; Daw, 2011; Valton et al., 2020). Each model was fit using 4 chains with 1,000 burn-in samples and 4,000 samples per chain except the test-retest models (including all sessions and an embedded correlation matrix between parameters) of the winning models which were fit using 2,000 samples per chain.

A number of visual and objective diagnostics of the MCMC performance were conducted to examine convergence of the model-fitting procedure (Ahn et al., 2017; Kruschke, 2015). Each model was inspected for divergences – of none were found for any of the models. All subject- and group-level parameters were checked for a Gelman-Rubin statistic ($\hat{R}$; Gelman & Rubin, 1992) value of less than 1.1 and an effective sample size (ESS) in the thousands. Trace plots of all subject- and group-level parameters were examined to ensure that the MCMC samples were well-mixed. If there were any issues with a model's MCMC performance, it was excluded as it would indicate that the model had not converged.

#### Four-armed bandit task

The restless four-armed bandit task assesses reward and punishment learning (Daw et al., 2006; Seymour et al., 2012). On each trial participants were asked to choose one out of four bandits (represented as boxes), which would display one out of four possible outcomes following a choice: reward (green token), punishment (red token), neither reward nor punishment (empty box) or both reward and punishment (red and green token). The probability of reward and punishment outcomes varied over time (with a slow random walk) independently of one another within each bandit, and independently between bandits. Varying reward and punishment probabilities over time, as opposed to using fixed probabilities, allows for prediction errors to be constantly produced over time, which improves the ability to estimate RL processes as participants are less likely to adopt a rule-based response policy, which can occur when probabilities are fixed. Participants were instructed on the non-stationary and independent nature of choice outcomes and were told that the goal was to maximize gains and minimize losses. The task lasted around 15 minutes with 200 trials in total.

##### Model-agnostic task analyses

Outcome measures for reliability analysis based on summary statistics:
p(stay) after win: number of repeated choices after wins only/total number of win trials
p(stay) after loss: number of repeated choices after losses only/total number of loss trials
p(stay) after neither: number of repeated choices after neither only/total number of neither trials

The probability to stay after wins, losses and neither were also measured using a mixed-effects logistic model for each session separately with random intercepts and slopes for participants using the lme4 package in R with the following model syntax:

stay~ 1+feedbackType+(1+ feedbackType |subjID),

where 'stay', the dependent variable, is a vector of 1 (choosing the same option as previous trial) or 0 (choosing a different option), and 'feedbackType', the predictor, is a vector coding for all task conditions per trial (previous trial feedback type: win, loss, neither or both). Reliability of p(stay) after a certain feedback derived from the mixed logistic regression is presented in Table S1.

| Mixed logistic regression | ICC(A,1) | ICC(1) | Pearson's r |
|---|---|---|---|
| Win | 0.46 (0.21-0.65) | 0.46 (0.21-0.65) | 0.46 (0.21-0.65) |
| Loss | 0.55 (0.33-0.72) | 0.55 (0.32-0.72) | 0.56 (0.33-0.73) |
| Neither | 0.67 (0.48-0.80) | 0.67 (0.48-0.80) | 0.67 (0.48-0.80) |

**Table S1: Reliability of model-agnostic p(stay) measures from hierarchical logistic regression models (conducted separately for each session) for the bandit task.** Brackets represent the 95% confidence interval.

#### Modelling results and diagnostics

Table S2 displays the seven models from the hBayesDM package fitted to the four-armed bandit data. The bandit4arm_lapse_decay model has previously shown to best fit the four-armed bandit task (Aylward et al., 2019). However, this model exhibited a number of Gelman−Rubin statistics $\hat{R}$ values greater than 1.1 and ESS <100. The ESS remained <100 for a number of parameters and trace plots showed poor mixing even when the sample size per chain was increased to 10,000 and default argument settings changed to: adapt_delta=0.99, stepsize=0.5, max_treedepth=20 (suggested parameter settings by hBayesDM defaults if model does not converge; Ahn et al., 2017). This indicates signs of poor convergence, and the bandit4arm_lapse_decay model was therefore excluded.

| Model | S1: LOOIC | S2: LOOIC |
|---|---|---|
| **Bandit4arm_lapse** | **21419.77** | **20469.63** |
| Bandit4arm_4par | 21426.74 | 20513.87 |
| Bandit4arm_singleA_lapse | 21663.92 | 20667.74 |
| Igt_pvl_decay | 21834.23 | 21060.78 |
| Igt_pvl_delta | 22392.60 | 21294.36 |
| Bandit4arm_2par_lapse | 25523.77 | 25421.07 |
| bandit4arm_lapse_decay* | - | - |

**Table S2: Model fits for the four-armed bandit task from the hBayesDM package.** The winning model has the lowest Leave-One-Out Information Criterion (LOOIC) and noted in bold here. S1: session 1; S2: session2. *This model exhibited poor convergence and was excluded.

The winning model, bandit4arm_lapse, was calculated by the following equations (Aylward et al., 2019):

$$Value_{t(i)}^{rew} = Value_{t(i)}^{rew} + \ Reward\ Learning\ Rate \times Prediction\ Error_{t(i)}^{rew}\ (1)$$

$$Value_{t(i)}^{pun} = Value_{t(i)}^{pun} + \ Punishment\ Learning\ Rate \times Prediction\ Error_{t(i)}^{pun}\ (2)$$

'Rew' and 'pun' refers to the reward (1,0) and punishment (0,-1) values on each trial (t) for a given bandit (i).

$$if\ i = chosen: Prediction\ Error_{t(i)}^{rew} = Reward\ Sensitivity\ \times\ Reward\ Outcome(t) - Value_{t-1(i)}^{rew}\ (3)$$

$$if\ i = unchosen: Prediction\ Error_{t(i)}^{rew} = -Value_{t-1(i)}^{rew}$$

$$if\ i = chosen: Prediction\ Error_{t(i)}^{pun} = Punishment\ Sensitivity\ \times\ Punishment\ Outcome(t) - Value_{t-1(i)}^{pun}\ (4)$$

$$if\ i = unchosen: Prediction\ Error_{t(i)}^{pun} = -Value_{t-1(i)}^{pun}$$

The subjective reward and punishment values were passed through a softmax function to estimate the probability of choosing a given bandit on each trial (j represents all bandits):

$$Choice\ Probability = \frac{\exp\left(Value_{t(i)}^{rew} + Value_{t(i)}^{pun}\right)}{\sum_j \exp\left(Value_{t(i)}^{rew} + Value_{t(i)}^{pun}\right)} \times (1 - Lapse) + \frac{Lapse}{4}\ (5)$$

In the RL model, the sensitivity parameters thus determine how much an outcome should be weighted ("how much an individual anticipates 'liking/disliking' an outcome"). In contrast, the learning rates determine how much weight should be given to the difference in expected and received outcomes ("how quickly an individual changes behaviour after punishments or sticks with a behaviour after rewards"). They thus describe related but different processes, and the parameters are uncorrelated in the current RL model (Figure S1). Incorporating both in the model allows for a deeper understanding of how people respond to outcomes. For example, in anhedonia individuals behave as if rewards are less rewarding but this could be due to either a lower sensitivity to rewards or slower learning of rewarding options. It is thus possible to examine whether symptoms might be associated with different learning rates or sensitivity (or both) in RL (compared with healthy individuals), which could inform different treatment strategies (Aylward et al., 2019). For example, if reward learning rates but not reward sensitivity is affected in depression – treatments should focus on altering how individuals integrate information over time rather than trying to increase sensitivity to rewarding outcomes. It is therefore useful to understand what the test-retest reliabilities are of these separate parameters.

To ensure that the winning model had identifiable parameters, we examined pair plots of the posterior distributions of the group-level parameters to determine that no major trade-off was occurring between parameters. Figure S1 displays corner plots of the group-level parameters in the Bandit4arm_lapse from session 1 and 2, exhibiting no major trade-offs between each other. Since the mean of the posterior distribution of parameters were used, we examined corner plots of all individuals' parameters to examine possible multimodal parameter estimates. Three individuals were excluded after identified multimodality, as the parameter point estimate (mean) would not be representative (Figure S2).

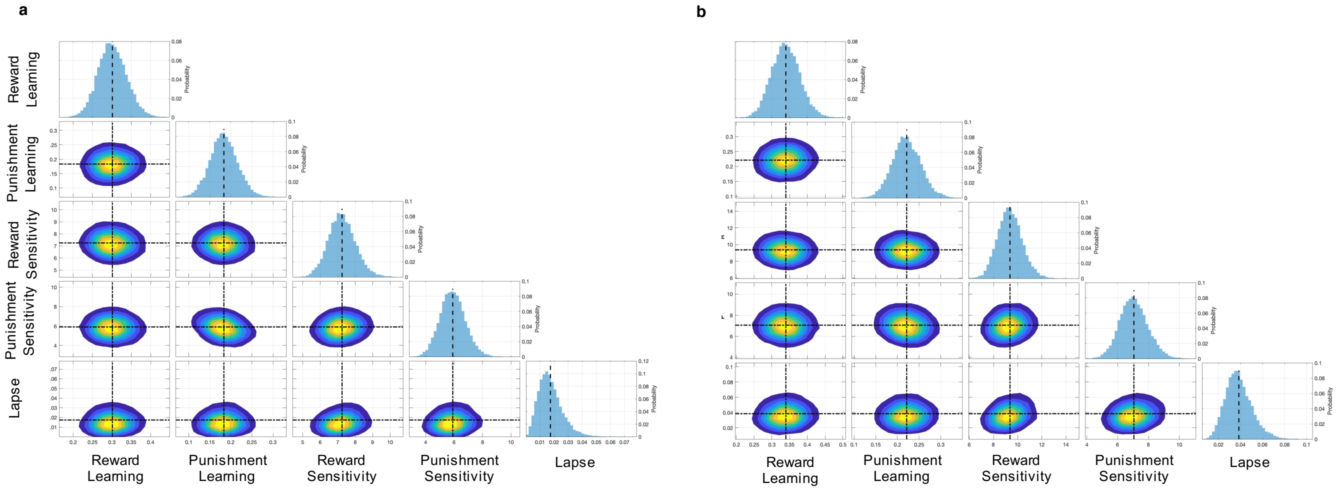


**Figure S1. Corner plots of all group-level parameters from the bandit4arm_lapse model**. The model was fitted to the four-armed bandit task, for session 1 (a) and session 2 (b). No parameters show major correlations. Dotted line indicates the mean.

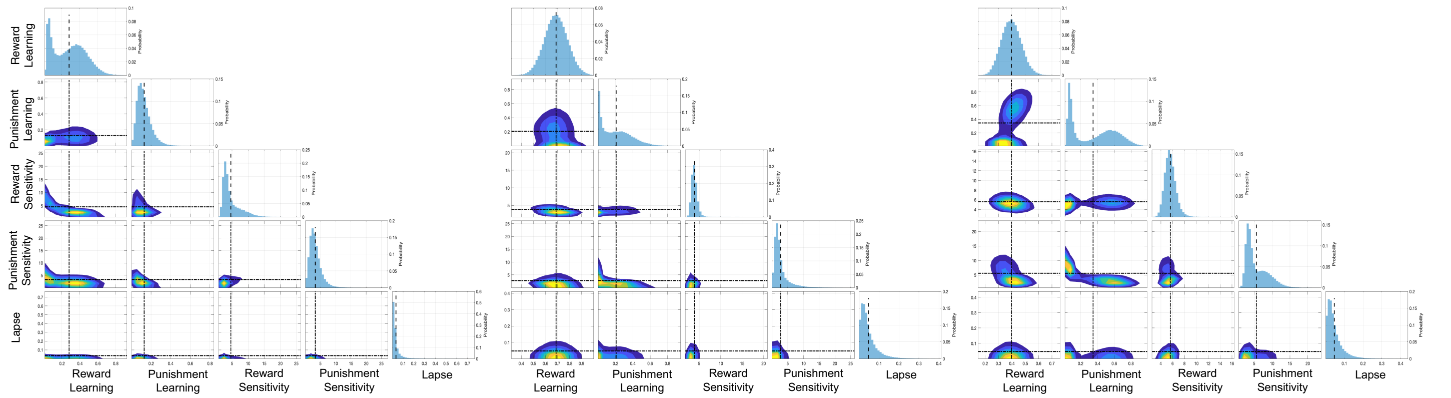


**Figure S2. Corner plots of parameters from the bandit4arm_lapse model for three individuals displaying multimodal parameter estimates.** Parameters from left to right and top to bottom are: 1) reward learning, 2) punishment learning, 3) reward sensitivity, 4) punishment sensitivity and 5) lapse.

Excluding these three individuals did not substantially affect the test-retest reliability.

We also examined if the winning model had recoverable parameters by generating simulated data for each participant based on their parameter estimates by adapting existing simulation scripts in R (Robinson, 2019). All parameters, except the lapse parameter, showed high recoverability (Figure S3). To examine if the winning model could also recapitulate real choices, simulated choices were generated for each individual using their estimated parameters and comparing the simulated choices with real choices (Figure S4).

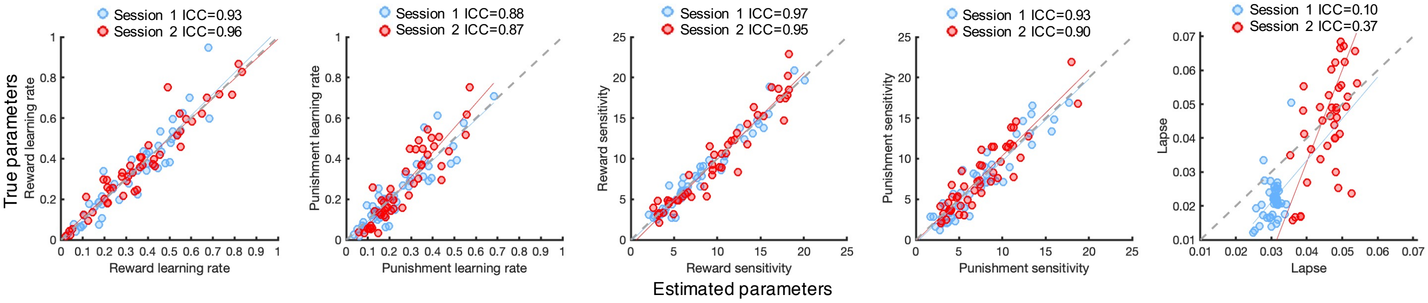


**Figure S3. Parameter recovery of the bandit4arm_lapse model.** Intraclass correlations (ICCs) represent two-way mixed model for single measures and absolute agreement.

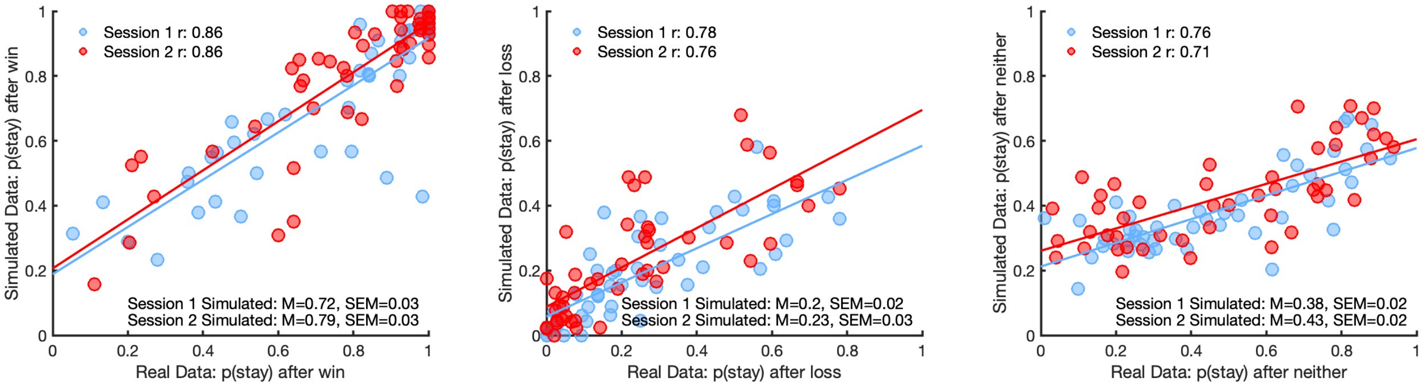


**Figure S4. Simulated versus real four-armed bandit data.** Simulated data based on participants' estimated parameters from the bandit4arm_lapse model, is plotted against the real data. R-values represent Pearson's correlations. P(stay): probability to stay; M: mean; SEM: standard error of the mean.

To better understand the poor recoverability of the lapse parameter, we also examined the standard deviations of the group-level parameters. The standard deviation of the lapse group-level parameter in session 1 exhibits a distribution with a positive skew, in contrast to session 2 where it takes on a bell-shaped curve instead (Figure S5). The parameter is also bounded at 0 with most of the samples close to 0. This likely accounts for the small variation in lapse parameter values as seen between individuals in session 1 (Figure S3; Figure 3).

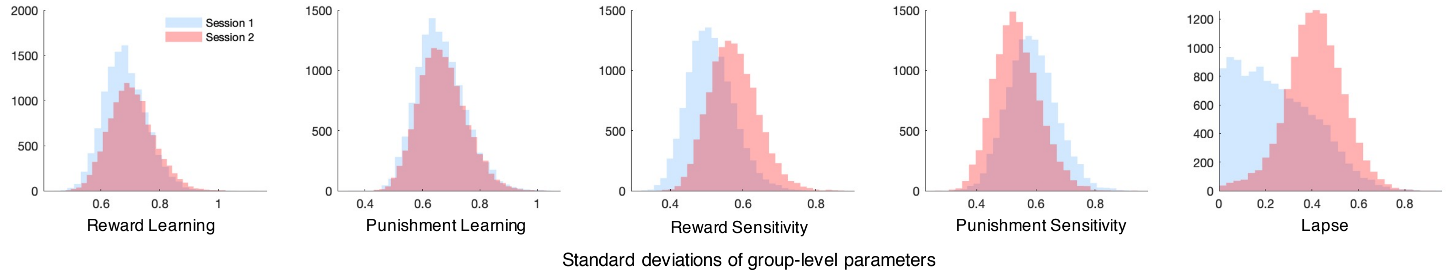


**Figure S5. Histograms of the standard deviations of all group-level parameters from the bandit4arm_lapse model for session 1 and session 2.**

Table S3 displays test-retest reliability and practice effects of Bandit4arm_lapse model parameters when estimated under a single hierarchical prior.

| Computational parameters | Test-retest reliability | | | Practice effects | | |
|---|---|---|---|---|---|---|
| **Four-armed bandit task (N=47)** | **ICC(A,1) (95% CI)** | **S1 Mean (SEM)** | **S2 Mean (SEM)** | **t** | ***p*** | ***$d_z$*** |
| Reward learning rate | 0.59 (0.37-0.75) | 0.34 (0.03) | 0.37 (0.03) | 1.00 | 0.32 | 0.14 |
| Punishment learning rate | 0.64 (0.43-0.78) | 0.24 (0.02) | 0.27 (0.03) | 1.42 | 0.16 | 0.20 |
| Reward sensitivity | 0.54 (0.31-0.72) | 8.48 (0.75) | 9.88 (0.80) | 1.91 | 0.06 | 0.27 |
| Punishment sensitivity | 0.45 (0.19-0.65) | 7.38 (0.69) | 7.76 (0.65) | 0.54 | 0.59 | 0.08 |
| Lapse | 0.02 (-0.27-0.31) | 0.04 (0.003) | 0.04 (0.003) | 0.29 | 0.77 | 0.04 |

**Table S3. Test-retest reliability and practice effects of all four-armed bandit task model parameters when estimated under a single prior (S1 and S2 data together).** CI: confidence interval; SEM: standard error of the mean; S1: session 1; S2: session 2.

#### Gambling task

The gambling task measures loss and risk aversion (Charpentier, Aylward, Roiser, & Robinson, 2017). On each trial, participants chose between a 50-50 gamble and a sure (guaranteed amount of points) option. The task was composed of two types of trials to disambiguate risk aversion from loss aversion. Loss aversion was measured using mixed-gamble trials, where the 50-50 gamble contained a gain and a loss, and the sure option 0 points. Risk aversion was assessed with gain-only trials, such that the 50-50 gamble resulted in either a gain or nothing and the sure option was a guaranteed gain.

An initial training phase was used to create individual calibrated offers in a second phase. The training phase used a staircase procedure to calibrate individual indifference points of loss/risk aversion (50 loss and 40 risk aversion trials). The second block contained 120 trials (64 loss and 56 risk aversion) centred on the individualized risk/loss aversion indifference points, which were presented in random order. Participants were instructed that there were two blocks of the task but not that the first block was a calibration phase. The task lasted 15 minutes.

##### Model-agnostic task analyses

Outcome measures for reliability analysis based on summary statistics:
P(gamble) on mixed trials: number of gamble choices/total number of mixed trials
P(gamble) on gain-only trials: number of gamble choices/total number of gain-only trials

The probability to gamble on mixed trials and gain-only trials were further measured using a mixed-effects logistic model for each session separately with random intercepts and slopes for participants using the following model syntax:

gamble~ 1+trialType+(1+ trialType |subjID),

where ‘gamble’, the dependent variable, is a vector of 1 (choosing the gamble option) or 0 (choosing the safe option), and ‘trialType’, the predictor, is a vector coding for trial type (mixed or gain-only trial). Reliability of p(gamble) depending on trial type derived from the mixed logistic regression is presented in Table S4.

| **Mixed logistic regression** | **ICC(A,1)** | **ICC(1)** | **Pearson's r** |
|---|---|---|---|
| Mixed trials | 0.63 (0.43-0.78) | 0.64 (0.44-0.78) | 0.63 (0.42-0.77) |
| Gain-only trials | 0.60 (0.39-0.75) | 0.60 (0.38-0.75) | 0.60 (0.39-0.76) |

**Table S4: Reliability of model-agnostic p(gamble) measures from hierarchical logistic regression models (conducted separately for each session) for the gambling task.** Brackets represent the 95% confidence interval.

#### Modelling results and diagnostics

Table S5 displays the three models from the hBayesDM package fitted to the gambling task data, where 1) ra_prospect includes a loss aversion, risk aversion and inverse temperature parameter, 2) ra_noLA includes a risk aversion and inverse temperature parameter, and 3) ra_noRA includes a loss aversion and inverse temperature parameter.

| **Model** | **S1: LOOIC** | **S2: LOOIC** |
|---|---|---|
| **Ra_prospect** | **5249.68** | **5160.39** |
| Ra_noLA | 6687.08 | 6551.38 |
| Ra_noRA | 6723.99 | 7167.25 |

**Table S5: Model fits for the gambling task from the hBayesDM package.** The winning model has the lowest Leave-One-Out Information Criterion (LOOIC) and noted in bold here. S1: Session 1; S2: Session 2.

The winning gambling task model, ra_prospect, was estimated with the following equations:

$$EV(gamble) = 0.5 \times gain(t)^{Risk\ aversion} + 0.5 \times Loss\ aversion \times -loss(t)^{Risk\ aversion} \quad (1)$$

$$EV(sure) = sure(t)^{Risk\ aversion} \quad (2)$$

On each trial (t) the subjective expected value (EV) of the gamble and sure option was calculated. These subjective expected values were passed through a softmax function to calculate the estimated probability of choosing the gamble option:

$$p(gamble) = \frac{1}{1 + \exp\left(-Inverse\ temperature \times [EV(gamble) - EV(sure)]\right)} \quad (3)$$

No major trade-offs between group-level parameters of the prospect model were identified (Figure S6). In addition, no individuals displayed multimodal parameter estimates.

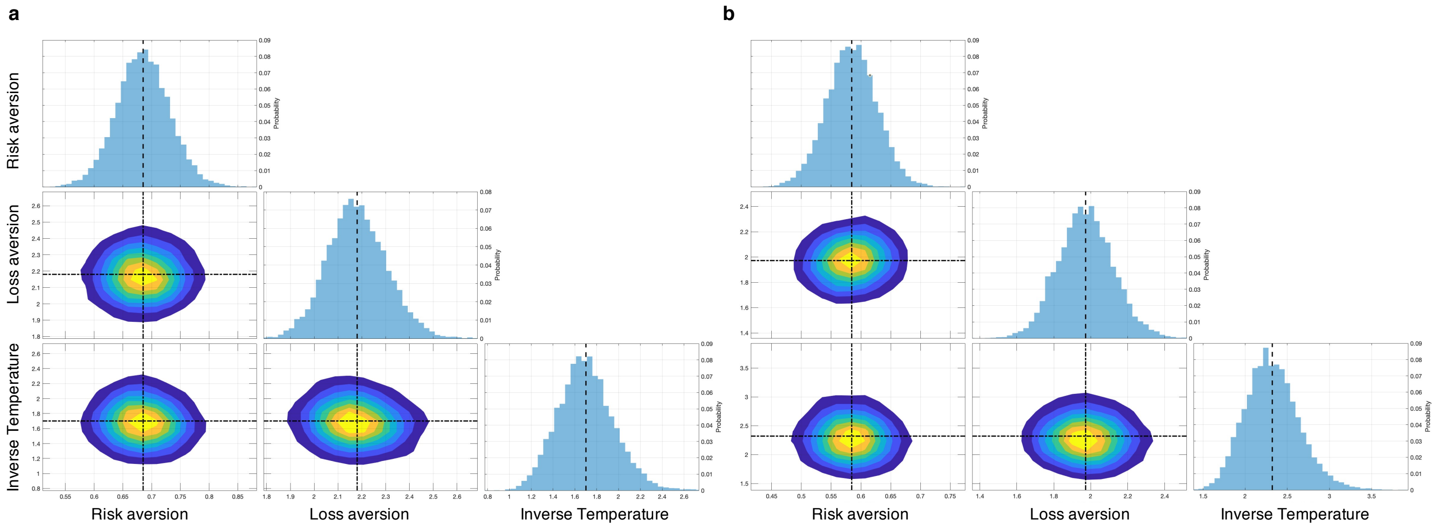


**Figure S6. Corner plots of all group-level parameters from the prospect theory model**. Models were fitted to the gambling task, for session 1 (a) and session 2 (b). No parameters show major correlations. Dotted line indicates the mean.

All parameters showed high recoverability (Figure S7). Simulated choices, generated from participants' estimated parameter values, tracked participants' real choices well (Figure S8).

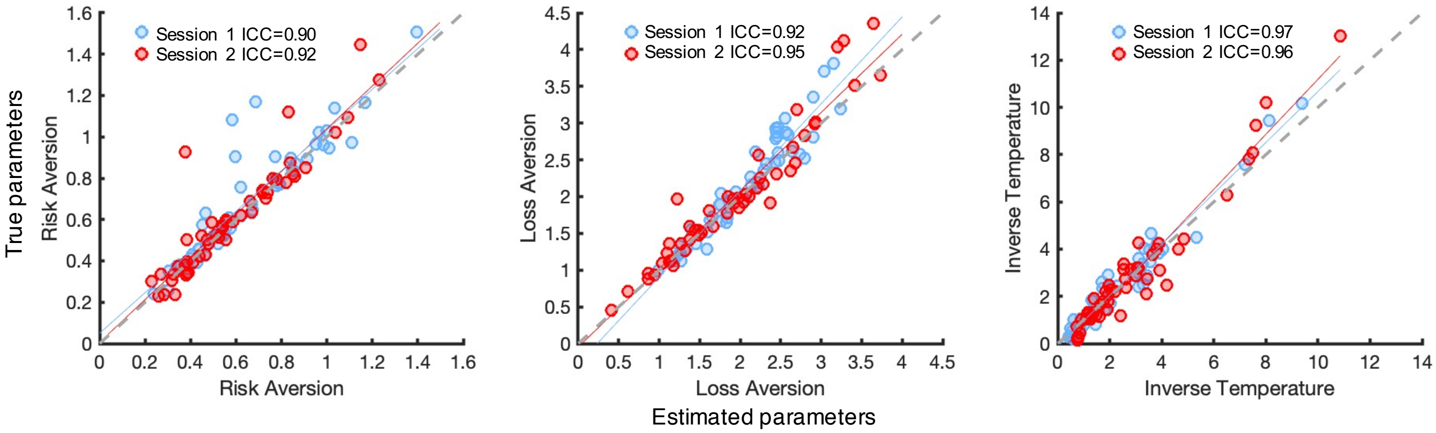


**Figure S7. Parameter recovery of the prospect model.** Intraclass correlations (ICCs) represent two-way mixed model for single measures and absolute agreement.

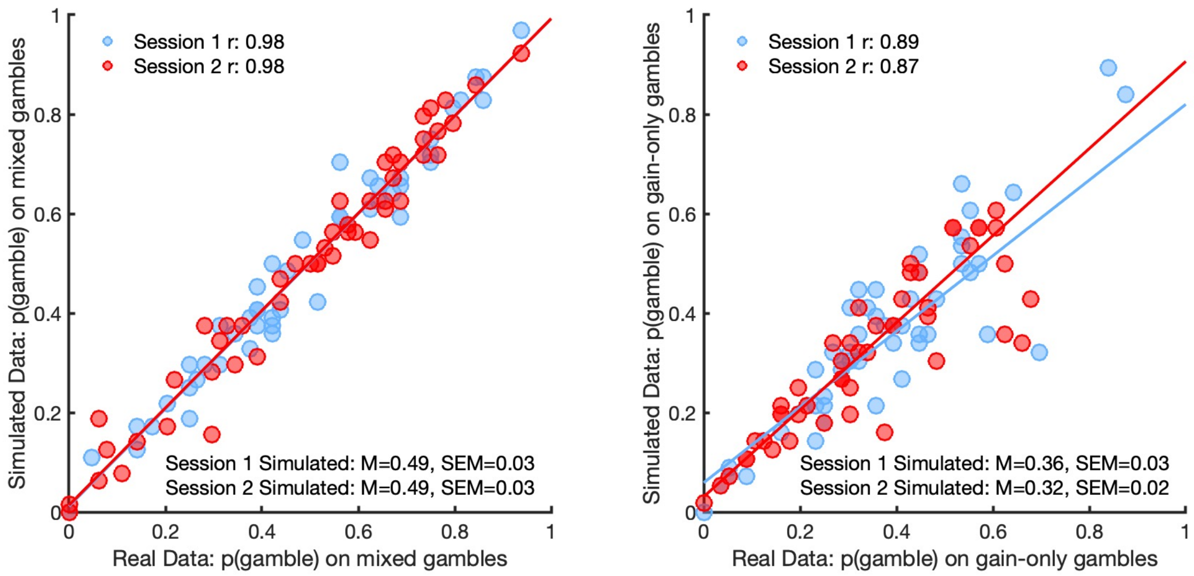


**Figure S8. Simulated versus real gambling task data.** Simulated gambling data, based on participants' estimated parameters from the prospect model, plotted against the real data. R-values represent Pearson's correlations. P(gamble): probability to gamble; M: mean; SEM: standard error of the mean.

Table S6 displays test-retest reliability and practice effects of ra_prospect model parameters when estimated under a single hierarchical prior.

| Computational parameters | Test-retest reliability | | Practice effects | | | |
|---|---|---|---|---|---|---|
| **Gambling task (N=49)** | **ICC(A,1) (95% CI)** | **S1 Mean (SEM)** | **S2 Mean (SEM)** | **t** | ***p*** | ***$d_z$*** |
| Loss Aversion | 0.70 (0.52-0.82) | 2.20 (0.11) | 2.01 (0.13) | 2.13 | 0.039 | 0.30 |
| Risk Aversion | 0.80 (0.60-0.89) | 0.70 (0.04) | 0.62 (0.04) | 3.62 | 0.001 | 0.52 |
| Inverse Temperature | 0.81 (0.65-0.89) | 2.40 (0.32) | 3.00 (0.39) | 2.88 | 0.006 | 0.41 |

**Table S6. Test-retest reliability and practice effects of all gambling task model parameters when estimated under a single prior (S1 and S2 data together).** ICC: intraclass correlation coefficient; CI: confidence interval; SEM: standard error of the mean; S1: session 1; S2: session 2.

#### Reliability calculations

The two-way mixed model for single measures and absolute agreement ICC (A,1) is defined as (Koo & Li, 2016; McGraw & Wong, 1996):

$$ICC = \frac{MS_B - MS_E}{MS_B + (k-1)MS_E + \frac{k}{n}(MS_W - MS_E)}$$

The one-way random model for single measures and absolute agreement ICC (1) is defined as (Koo & Li, 2016; McGraw & Wong, 1996):

$$ICC = \frac{MS_B - MS_R}{MS_B + (k-1)MS_R},$$

where $MS_B$= mean square of between-subject variance, $MS_E$=mean square of the error variance, $MS_W$=mean square of the within-subject variance, $MS_R$= mean square for residual sources of variance, *k*=number of testing sessions, *n*= number of subjects.

#### Exploratory demographic correlations

To explore if any demographic characteristics may have a relationship with any of the task measures, Pearson's correlations were used to correlate task measures (model-agnostic outcomes and parameter estimates) with age and years of education. Independent t-tests were used to examine the effect of gender on task measures. There were no significant associations between any of the task measures and demographic measures when correcting for multiple comparisons (Table S7).

| **Measures** | **Age** | | **Years of education** | | **Gender** | |
|---|---|---|---|---|---|---|
| | **r** | ***p*** | **r** | ***p*** | **t** | ***p*** |
| **Four-armed bandit task** | | | | | | |
| *Model-agnostic summary statistics (N=50)* | | | | | | |
| p(stay) loss – p(stay) neither | -0.23 | 0.10 | -0.01 | 0.95 | 0.83 | 0.41 |
| p(stay) win – p(stay) neither | 0.01 | 0.95 | -0.05 | 0.72 | 0.18 | 0.86 |
| *Computational parameters (N=47)* | | | | | | |
| Reward learning rate | -0.09 | 0.57 | -0.18 | 0.22 | 0.44 | 0.67 |
| Punishment learning rate | 0.33 | 0.03 | 0.14 | 0.34 | 0.70 | 0.49 |
| Reward sensitivity | 0.12 | 0.43 | 0.26 | 0.08 | 0.74 | 0.46 |
| Punishment sensitivity | 0.06 | 0.68 | 0.06 | 0.68 | 0.97 | 0.34 |
| Lapse | -0.004 | 0.98 | -0.11 | 0.47 | 1.04 | 0.30 |
| **Gambling task** | | | | | | |
| *Model-agnostic summary statistics (N=49)* | | | | | | |
| p(gamble) mixed trials | 0.11 | 0.44 | 0.20 | 0.17 | 0.61 | 0.55 |
| p(gamble) gain-only trials | 0.05 | 0.74 | 0.13 | 0.37 | 0.04 | 0.97 |
| *Computational parameters (N=49)* | | | | | | |
| Loss Aversion | -0.12 | 0.41 | -0.01 | 0.50 | 1.30 | 0.20 |
| Risk Aversion | 0.05 | 0.71 | 0.23 | 0.12 | 1.09 | 0.28 |
| Inverse Temperature | 0.10 | 0.51 | 0.06 | 0.70 | 1.02 | 0.32 |

**Table S7. Associations between demographic and task measures.** R-values represent Pearson's correlations and t-values represent independent t-tests. Raw p-values are presented. Correcting for multiple comparisons results in an alpha level of 0.05/12=0.004.

#### Supplementary Discussion

##### *Practice effects*

Some of the reliable parameters showed small-to-medium session effects. Practice effects can either obscure a true effect or lead to false treatment claims if appropriate controls are not employed. Quantifying session effects, however, allows such changes to be accounted for. In the RL model, reward sensitivity increased in the second session, while all other reliable RL processes were fairly stable. In contrast, all PT parameters showed significant session effects. However, the sensitivity analysis showed no substantial practice effects of RL parameters when data were fit under a single prior, while the practice effects of PT parameters remained. Thus, the apparent practice effects on the RL task should be interpreted with caution, as these may be overestimated under the two prior estimation approach (Valton et al., 2020). Importantly, the ICC estimates were hardly influenced by the estimation approach.
